# Obstacles to inferring mechanistic similarity using Representational Similarity Analysis

**DOI:** 10.1101/2022.04.05.487135

**Authors:** Marin Dujmović, Jeffrey S Bowers, Federico Adolfi, Gaurav Malhotra

## Abstract

Representational Similarity Analysis (RSA) is an innovative approach used to compare neural representations across individuals, species and computational models. Despite its popularity within neuroscience, psychology and artificial intelligence, this approach has led to difficult-to-reconcile and contradictory findings, particularly when comparing primate visual representations with deep neural networks (DNNs). Here, we demonstrate how such contradictory findings could arise due to incorrect inferences about mechanism when comparing complex systems processing high-dimensional stimuli. In a series of studies comparing computational models, primate cortex and human cortex we find two problematic phenomena: a “mimic effect”, where confounds in stimuli can lead to high RSA-scores between provably dissimilar systems, and a “modulation effect”, where RSA- scores become dependent on stimuli used for testing. Since our results bear on a number of influential findings, we provide recommendations to avoid these pitfalls and sketch a way forward to a more solid science of representation in cognitive systems.

## Introduction

How do other animals see the world? Do different species represent the world in a similar manner? How do the internal representations of AI systems compare with humans and animals? The traditional scientific method of probing internal representations of humans and animals (popular in both psychology and neuroscience) relates them to properties of the external world. By moving a line across the visual field of a cat, Hubel & Wisel [1] found out that neurons in the visual cortex represent edges moving in specific directions. In another Nobel-prize winning work, O’Keefe, Moser & Moser [2, 3] discovered that neu-rons in the hippocampus and entorhinal cortex represent the location of an animal in the external world. Despite these successes it has proved difficult to relate internal repre-sentations to more complex properties of the world. Moreover, relating representations across individuals and species is challenging due to the differences in experience across individuals and differences of neural architectures across species.

These challenges have led to recent excitement around Representation Similarity Analysis (RSA), which is a multi-voxel pattern analysis method specifically designed to compare representations between different systems. RSA usually takes patterns of activity from two systems and computes how the distances between activations in one system correlate with the distances between corresponding activations in the second system (see Fig 1). Rather than compare each pattern of activation in the first system directly to the corresponding pattern of activation in the second system, it computes representational distance matrices (RDMs), a *second-order* measure of similarity that compares systems based on the relative distances between neural response patterns. This arrangement of neural response patterns in a representational space has been called a system’s *represen-tational geometry* [4]. The advantage of looking at representational geometries is that one no longer needs to match the architecture of two systems, or even the feature space of the two activity patterns. One could compare, for example, fMRI signals with single cell recordings, EEG traces with behavioural data, or vectors in a computer algorithm with spiking activity of neurons [5]. RSA is now ubiquitous in computational psychology and neuroscience and has been applied to compare object representations in humans and primates [6], representations of visual scenes by different individuals [7,8], representations of visual scenes in different parts of the brain [9], to study specific processes such as cog-nitive control [10] or the dynamics of object processing [11], and most recently, to relate neuronal activations in human (and primate) visual cortex with activations of units in Deep Neural Networks [12–16].

**Figure 1:**
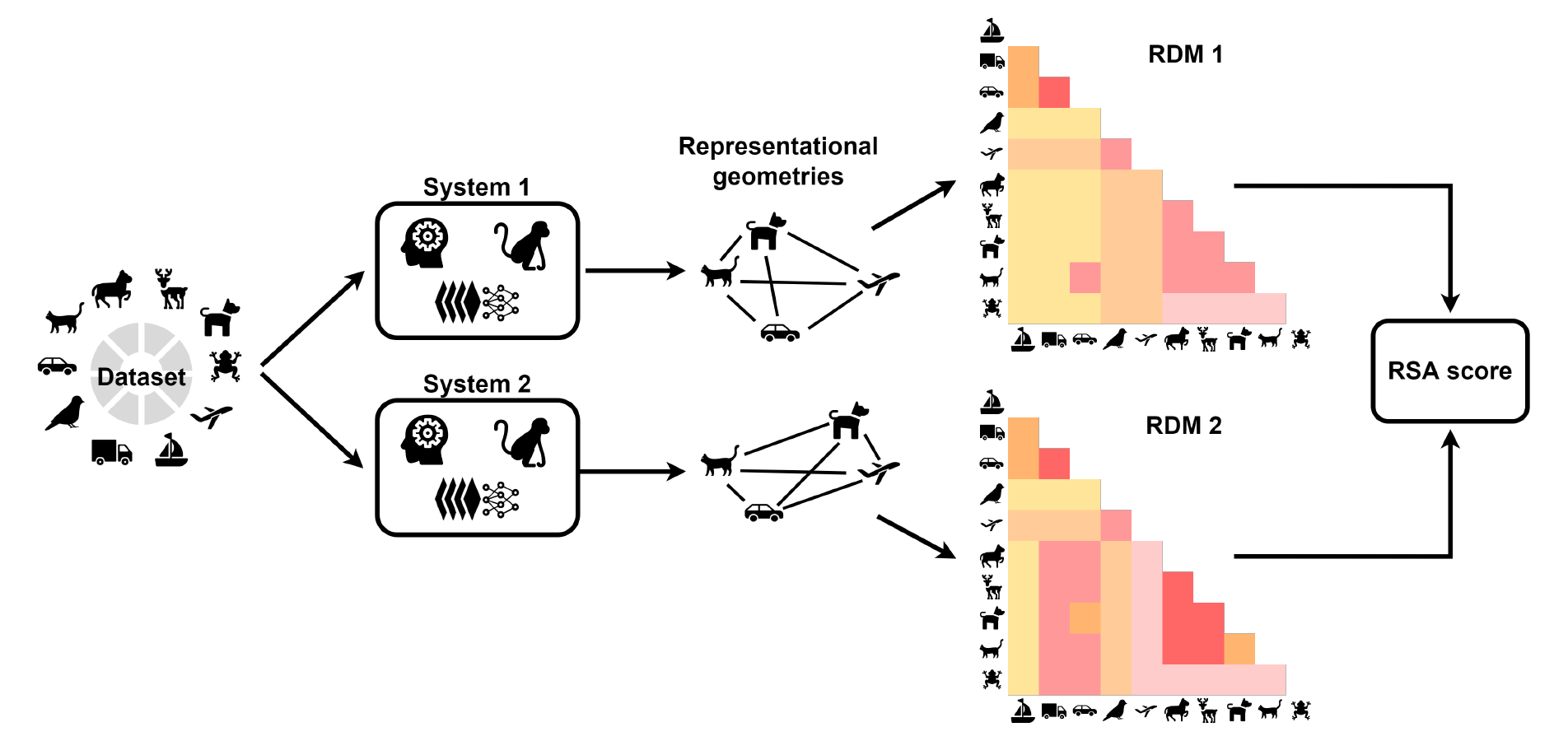
RSA calculation. Stimuli from a set of categories (or conditions) are used as inputs to two different systems (for example, a human brain and a primate brain). Activity from regions of interest is recorded for each stimulus. Pair-wise distances in activity patterns are calculated to get the representational geometry of each system. This representational geometry is expressed as a representational dissimilarity matrix (RDM) for each system. Finally, an RSA score is determined by computing the correlation between the two RDMs. It is up to the resercher to make a number of choices during this process including the choice of distance measure (e.g., 1-Pearson’s r, Euclidean distance etc.) and a measure for comparing RDMs (e.g., Pearson’s *r*, Spearman’s *ρ*, Kendall’s *τ*, etc.).

However, this flexibility in the application of RSA comes at the price of increased ambiguity in the inferences one can draw from this analysis. Since RSA is a second-order statistic (it looks at the similarity of similarities), it remains ambiguous which stimulus features drive the observed representational geometry in each system [17]. That is, two systems that operate on completely different stimulus features can nevertheless have highly correlated representational geometries. This makes inferences about mech-anism based on conducting RSA highly problematic. While researchers have recently highlighted a similar conceptual issue of confounds driving performance for multivariate decoding methods [18–20], it is less well appreciated for RSA. This is likely because there is a lack of understanding of how confounds can lead to misleading RSA scores, whether it is plausible that such confounds exist in datasets used for RSA and whether existing methods of dealing with confounds can address problematic inferences.

In our view, one particularly problematic area is research comparing biological systems and Deep Neural Networks (DNNs). There are many examples in this domain where researchers have recently used RSA for making inferences about psychological and neural mechanisms. For example, Cichy et al. [15] observed a correspondence in the RDMs of DNNs performing object categorization and neural responses in human visual cortex recorded using MEG and fMRI. Based on this correspondence, the authors concluded that:

> …hierarchical systems of visual representations emerge in both the human ventral and dorsal visual stream *as the result of* task constraints of object categorization posed in everyday life, and provide strong evidence for object representations in the dorsal stream independent of attention or motor intention. [pg. 5, emphasis added]

Thus, the correspondence in RDMs is used to infer the mechanism of emergence of visual representations. Based on a similar comparison, Kriegeskorte [21] concluded that ^1^:

> Deep convolutional feedforward networks for object recognition are not biologically detailed and rely on nonlinearities and learning algorithms that may differ from those of biological brains. Nevertheless they learn internal repre-sentations that are highly similar to representations in human and nonhuman primate IT cortex. [pg. 441]

In this paper, we will show – through a series of simulations – that such inferences about similarity in neural representations or mechanism based on RSA are unwarranted. We will show that when target RDMs are obtained using a highly complex model and high-dimensional stimuli (both of which are true for comparisons between biological systems and DNNs) unknown confounds could drive similarity in representational geometries. Under these conditions, methods developed for dealing with confounds in multivariate decoding, such as cross-validation [23] and confound regression [24] may be insufficient for preventing false inferences. Furthermore, we will show that these confounds are not just possible, but also plausible given the nature of stimuli and structure of datasets frequently used for testing similarities between DNNs and humans. We will also argue that representational geometry should *not* be understood as the representation of a system as it conflicts with how most psychologists and neuroscientists view representations – the relationship between cognitive states and entities in the external world (see section S1 of the Supplementary Infomration a brief history of RSA and its philosophical origins).

The structure of the paper is as follows. In Study 1, in a bare-bones setup, we show that it is possible for two systems to transform input stimuli through known functions that are vastly different but end up with similar representational geometries. In particular, the study shows that 1) the presence of second-order confounds in the training data can lead systems to mimic each other’s representational geometry even in the absence of mechanistic similarity, and 2) the intrinsic structure of datasets rather than mechanistic alignment can lead to artifactual modulation of RSA scores. Then in Studies 2 and 3 we use real neural data collected in previous experiments to show these problems extend to more complex datasets directly relevant to artificial intelligence and computational neuroscience. Finally, in Study 4, we show that not only are misleadingly high RSA scores possible in practice but they are also highly plausible given the hierarchical structure of categories in datasets that are routinely used. Since our results have considerable generality with respect to current practices across multiple fields, we discuss the implications for published results, including a discussion of two alternative philosophical perspectives on the nature of mental representations that our findings speak to. We conclude by provid-ing some general recommendations regarding how to best compare representations across systems going forward.

## Results

### Proof of concept

It may be tempting to infer that two systems which have similar representational geometries for a set of concepts do so because they encode similar properties of sensory data and transform sensory data through a similar set of functions. In this section, we show that it is possible, at least in principle, for qualitatively different systems to end up with very similar representational geometries even though they (i) transform their inputs through very different functions, and (ii) select different features of inputs.

### Study 1: Demonstrably different transformations of inputs can lead to low or high RSA-scores

We start by considering a simple two-dimensional dataset and two systems where we know the closed-form functions that project this data into two representational spaces. This simple setup helps us gain a theoretical understanding of the circumstances under which it is possible for qualitatively different projections to show similar representational geometries.

Consider a population of animate and inanimate objects that consist of four categories of objects – birds, dogs, airplanes and bicycles. Each object in this population will have a set of stimulus features, using which one can map each exemplar from all four categories into a feature space. In Fig 2A (left), we show a hypothetical 2D feature space where exemplars from each category cluster together. Futhermore, we consider two datasets sampled from this population – Dataset A (Fig 2A, middle) which consists of birds and bicycles and Dataset B (Fig 2A, right) which consists of dogs and airplanes. Both datasets consist of animate and inanimate objects, but they differ in how items in each category are represented in the input space.

**Figure 2:**
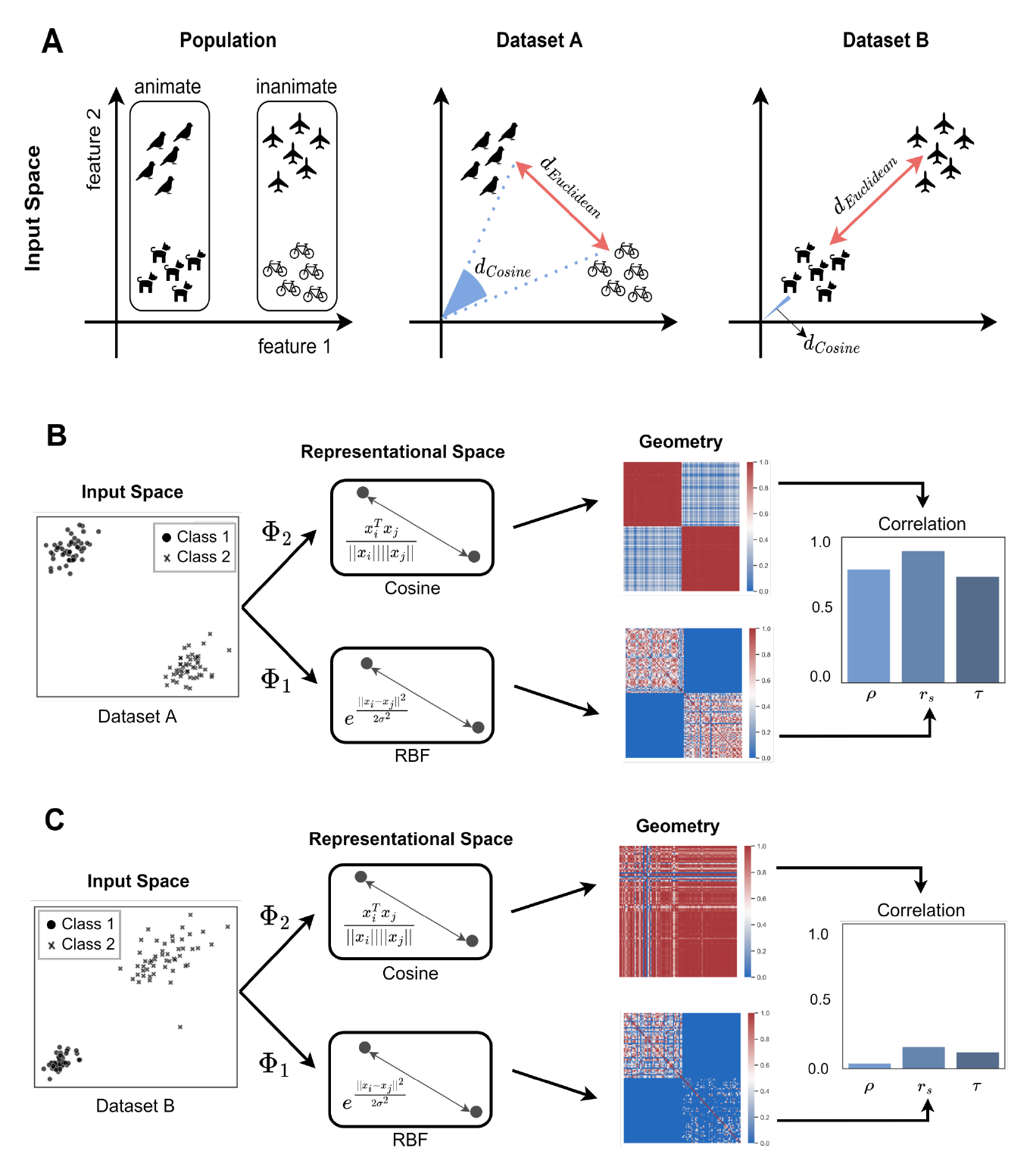
Mimic and modulation effect in representational geometries. (A) An example of a population of animate (birds, dogs) and inanimate (planes, bikes) objects, plotted in a hypothetical 2D stimulus feature space. Two datasets are sampled from this population: In Dataset A (middle), the Euclidean distance (in input space) between categories mirrors the Cosine distance, while in Dataset B (right) it does not. (B) Simulation where two systems transform stimuli in Dataset A into latent represen-tations such that the (dot product) distance between latent vectors is given by RBF and Cosine kernels, respectively. As Euclidean and Cosine distances in the input space mirror each other, the representational geometries (visualised here using kernel matrices) end up being highly correlated (shown using Pearson (*ρ*), Spearman (*r_s_*) and Kendall’s (*τ*) correlation coefficients on the right). We call this strong correlation in representational geometries despite a difference in input transformation a *mimic effect*. (C) Simulation where objects in Dataset B are projected using same transformations as (B). The (dot product) distance is still given by the same (RBF and Cosine) kernels. However, for this dataset, the Euclidean and Cosine distances in input space do *not* mirror each other and as a consequence, the representational geometries show low correlation. Thus the correlation in representational geometries depends on how the datasets are sampled from the population. We call this change in correlation a *modulation effect*.

Now, consider two information-processing systems that re-represent Dataset A into two different latent spaces (Fig 2B). These could be two recognition systems designed to distinguish animate and inanimate categories. We assume that we can observe the repre-sentational geometry of the latent representations of each system and we are interested in understanding whether observing a strong correlation between these geometries implies whether the two systems have a similar *representational space* – that is, they project in-puts into the latent space using similar functions. To examine this question, we consider a setup where we know the functions, Φ_1_ and Φ_2_, that map the inputs to the latent space in each system. We will now demonstrate that even when these functions are qualitatively different from each other, the geometry of latent representations can nevertheless be highly correlated. We will also show that the difference in representational spaces becomes more clear when one considers a different dataset (Dataset B), where inputs projected using the same functions now lead to a low correlation in representational geometries.

We can compute the geometry of a set of representations by establishing the pair-wise distance between all vectors in each representational space Φ. There are many different methods of computing this representational distance between any pair of vectors, all de-riving from the dot product between vectors (see, for example, Fig 1 in [25]). Previous research has shown that the choice of the distance metric itself can influence the inferences one can draw from one’s analysis [25,26]. However, here our focus is not the distance met-ric itself, but the fundamental nature of RSA. Therefore, we use the same generic distance metric – the dot product – to compute the pair-wise distance between all vectors in both representational spaces. In other words, the representational distance *d*[Φ(***x_i_***), Φ(***x_j_*)**], between the projections of any pair of input stimuli, ***x_i_*** and ***x_j_*** into a feature space Φ, is proportional to the inner product between the projections in the feature space:

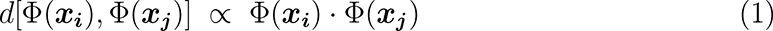

And we can obtain the representational geometry of the input stimuli {***x*_1_**,…, ***x_n_***} in any representational space Φ by computing the pairwise distances, *d*[Φ(***x_i_***), Φ(***x_j_***)] for all pairs of data points, (*i, j*). Here, we assume that the projections Φ_1_ and Φ_2_ are such that these pairwise distances are given by two positive semi-definite kernel functions *κ*_1_(***x_i_, x_j_***) and *κ*_2_(***x_i_, x_j_***), respectively:

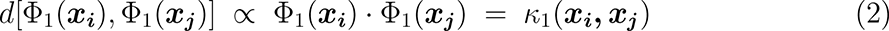

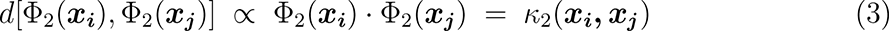

Now, let us consider two qualitatively different kernel functions: 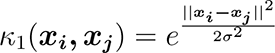 is a radial-basis kernel (where *σ*^2^ is the bandwidth parameter of the kernel), while *κ*_2_(***x_i_, x_j_***) 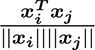 is a cosine kernel. We have chosen Φ_1_ and Φ_2_ such that they are two fundamen-tally different projections of the inputs {***x*_1_**,…, ***x_n_***} – while Φ_2_ maps a 2D input ***x_i_*** into a 2D feature space, Φ_1_ maps the same 2D input into an infinite-dimensional space. Nev-ertheless, since cosine and RBF kernels are Mercer kernels, we can compute the distances (as measured by the dot product) between each pair of projected vectors using the kernel trick [27,28]. That is, we can find the distance between any pair of points in the represen-tational space by applying the kernel function to those points in the input space. These pairwise distances are shown by the kernel matrices in Fig 2B.

Next, we can determine how the geometry of these projections in the two systems relate to each other by computing the correlation between the kernel matrices, shown on the right-hand-side of Fig 2B. We can see from these results that the kernel matrices are highly correlated – i.e., the input stimuli are projected to very similar geometries in the two representational spaces.

If one did not know the input transformations and simply observed the correlation between kernel matrices, it would be tempting to infer that the two systems Φ_1_ and Φ_2_ transform an unknown input stimulus ***x*** through a similar set of functions – for example functions that belong to the same class or project inputs to similar representational spaces. However, this would be an error. The projections Φ_1_(***x***) and Φ_2_(***x***) are fundamentally different – Φ_1_ (radial basis kernel) projects an input vector into an infinite dimensional space, while Φ_2_ (cosine kernel) projects it onto a unit sphere. The difference between these functions becomes apparent if one considers how this correlation changes if one considers a different set of input stimuli. For example, the set of data points from Dataset B (sampled fromt the same population) are projected to very different geometries, leading to a low correlation between the two kernel matrices (Fig 2C).

In fact, the reason for highly correlated kernel matrices in Fig 2B is not a similarity in the transformations Φ_1_ and Φ_2_ but the structure of the dataset. The representational distance between any two points in the first representational space, *d*[Φ_1_(***x_i_***), Φ_1_(***x_j_***)], is 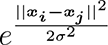. That is, the representational distance in Φ_1_ is a function of their Euclidean distance *||****x_i_ − x_j_****||* in the input space. On the other hand, the representational distance between any two points in the second representational space, *d*[Φ_2_(***x_i_***), Φ_2_(***x_j_***)], is, 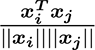. That is, the representational distance in Φ_2_ is a function of their cosine distance in the input space. These two stimulus features – Euclidean distance and cosine distance – are *confounds* that lead to the same representational geometries for certain datasets. In Dataset A, the stimuli is clustered such that the Euclidean distance between any two stimuli is correlated with their cosine distance (see Fig 2A, middle). However, for Dataset B, the Euclidean distance is no longer correlated with the angle (see Fig 2A, right) and the confounds lead to different representational geometries, as can be seen in Fig 2C. Thus, this example illustrates two effects: (i) a *mimic* effect, where two systems that transform sensory input through very different functions end up with similar representational geometries (Fig 2B), and (ii) a *modulation* effect, where two systems that are non-identical have similar representational geometries for one set of inputs, but dissimilar geometries for a second set (compare Fig 2B and 2C).

### Study 2: Complex systems encoding different features of inputs can show a high RSA-score

Study 1 made a number of simplifying assumptions – the dataset was two-dimensional, clustered into two categories and we intentionally chose functions Φ_1_ and Φ_2_ such that the kernel matrices were correlated in one case and not correlated in the other. It could be argued that, even though the above results hold in principle, they are unlikely in practice when the transformations and data structure are more complex. For example, it might be tempting to assume that accidental similarity in representational geometries becomes less likely as one increases the number of categories (i.e., clusters or conditions) being considered. However, In Fig 3 we illustrate how complex systems transforming high-dimensional input from a number of categories may achieve high RSA scores. Even though one system extracts surface reflectance and the other extracts global shape, they can end up with very similar representational geometries. This would occur if objects similar in their reflectance properties were also similar in shape (e.g., glossy balloons and light bulbs) and if objects dissimilar according to reflectance properties were also dissimilar in shape (e.g., dogs and light bulbs). This is the mimic effect, where representational geometries of these two systems end up being similar because reflectance and shape are second-order confounds in this dataset. Conducting RSA on this dataset will show a high correlation in RDMs, even though the latent representations in these systems are related to very different stimulus features.

**Figure 3:**
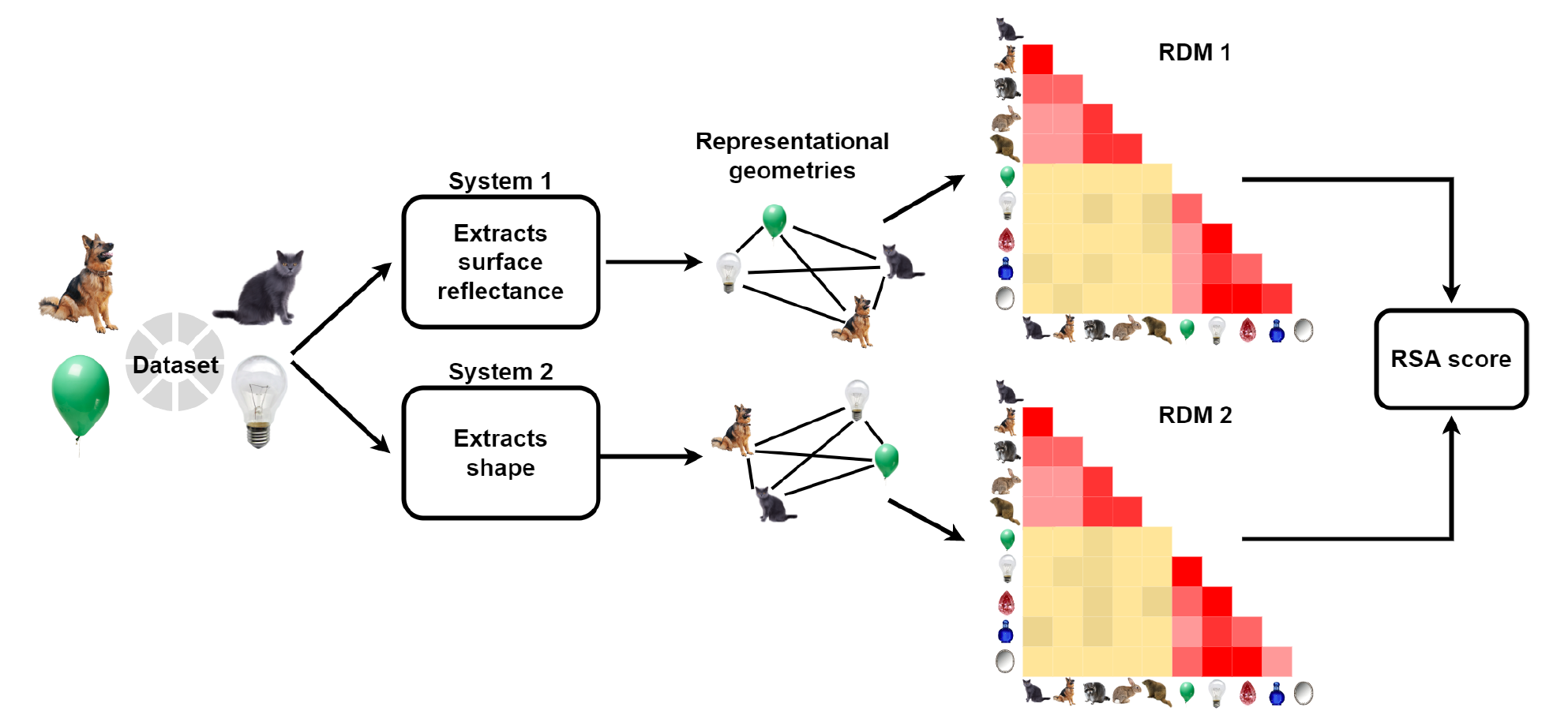
Example of a second-order confound. Two systems, one forming representations based on surface reflectance of objects (while ignoring all other features such as colour or texture) and the other based on global shape (while ignoring other features), can have very similar representational geometries. This similarity would lead to a high RSA score but would not justify an inference about the representations being similar.

To demonstrate this empirically, we now consider a more complex setup, where the transformations Φ_1_ and Φ_2_ are modelled as feedforward deep neural networks (DNNs), trained to classify a high-dimensional dataset into multiple categories. Many studies that use RSA compare systems using naturalistic images as visual inputs [6, 12]. While using naturalistic images brings research closer to the real-world, it is also well-known that datasets of naturalistic images frequently contain confounds – independent features that can predict image categories [29]. We will now show how the simplest of such confounds, a single pixel, can lead to a high RSA score between two DNNs that encode qualitatively different features of inputs.

Consider the same setup as above, where an input stimulus, ***x***, is transformed to a representation space by two systems, Φ_1_ and Φ_2_. Instead of a two-dimensional input space, ***x*** now exists in a high-dimensional image space and Φ_1_ and Φ_2_ are two versions of a DNN – `VGG-16` – trained to classify input images into different categories. We ensured that Φ_1_ and Φ_2_ were qualitatively different transformations of input stimuli by making the networks sensitive to different predictive features within the stimuli. The first network was trained on an unperturbed dataset, while the second network was trained on a modified version of the dataset, where each image was modified to contain a confound – a single pixel in a location that was diagnostic of the category (see Fig 4 for the general approach).

**Figure 4:**
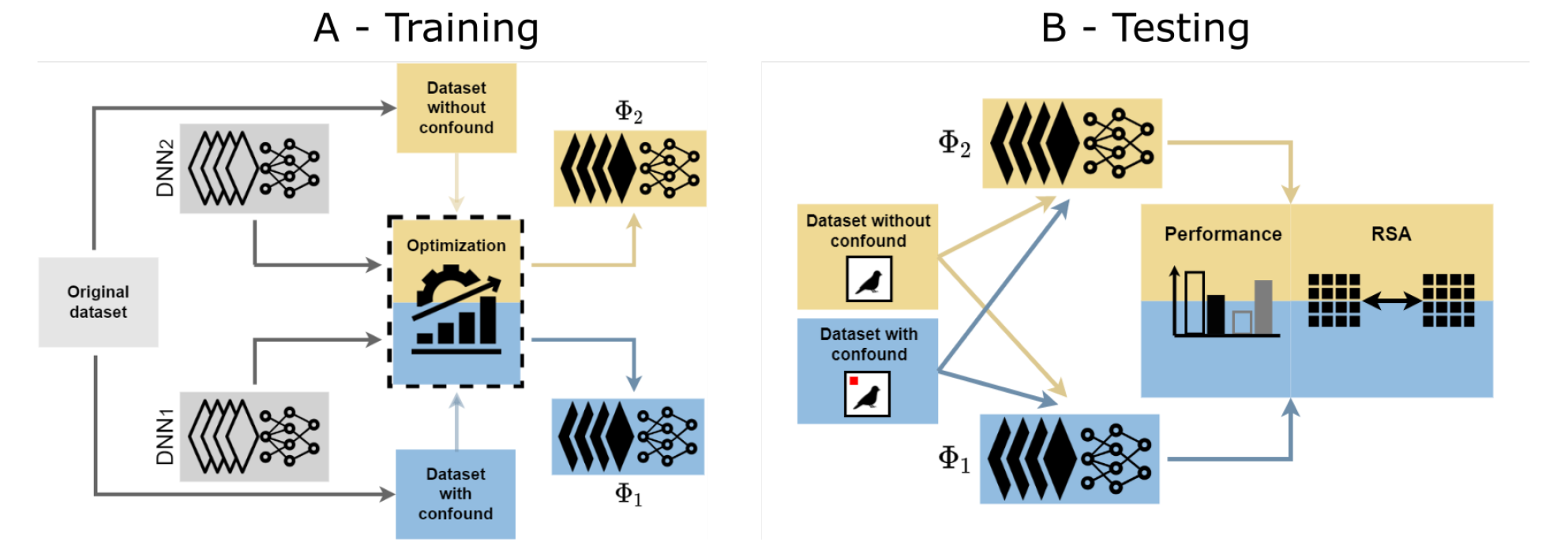
Training and testing DNNs with different feature encodings. Panel A shows the training procedure for Studies 2–4, where we created two versions of the original dataset (gray), one containing a confound (blue) and the other left unperturbed (yellow). These two datasets were used to train two networks (gray) on a categorisation task, resulting in two networks that learn to categorise images either based on the confound (projection Φ_2_) or based on statistical properties of the unperturbed image (projection Φ_1_). Panel B shows the testing procedure where each network was tested on stimuli from each dataset – leading to a 2×2 design. Performance on these datasets was used to infer the features that each network encoded and their internal response patterns were used to calculate RSA-scores between the two networks.

The locations of these diagnostic pixels were chosen such that they were correlated to the corresponding representational distances between classes in Φ_1_. Our hypothesis was that if the representational distances in Φ_2_ preserve the physical distances of diagnostic pixels in input space, then this confound will end up mimicking the representational geometry of Φ_1_, even though the two systems use qualitatively different features for classification. Furthermore, we trained two more networks, Φ_3_ and Φ_4_, which were identical to Φ_2_, except these networks were trained on datasets where the location of the confound was uncorrelated (Φ_3_) or negatively correlated (Φ_4_) with the representational distances in Φ_1_ (see Fig 5 and Methods for details).

**Figure 5:**
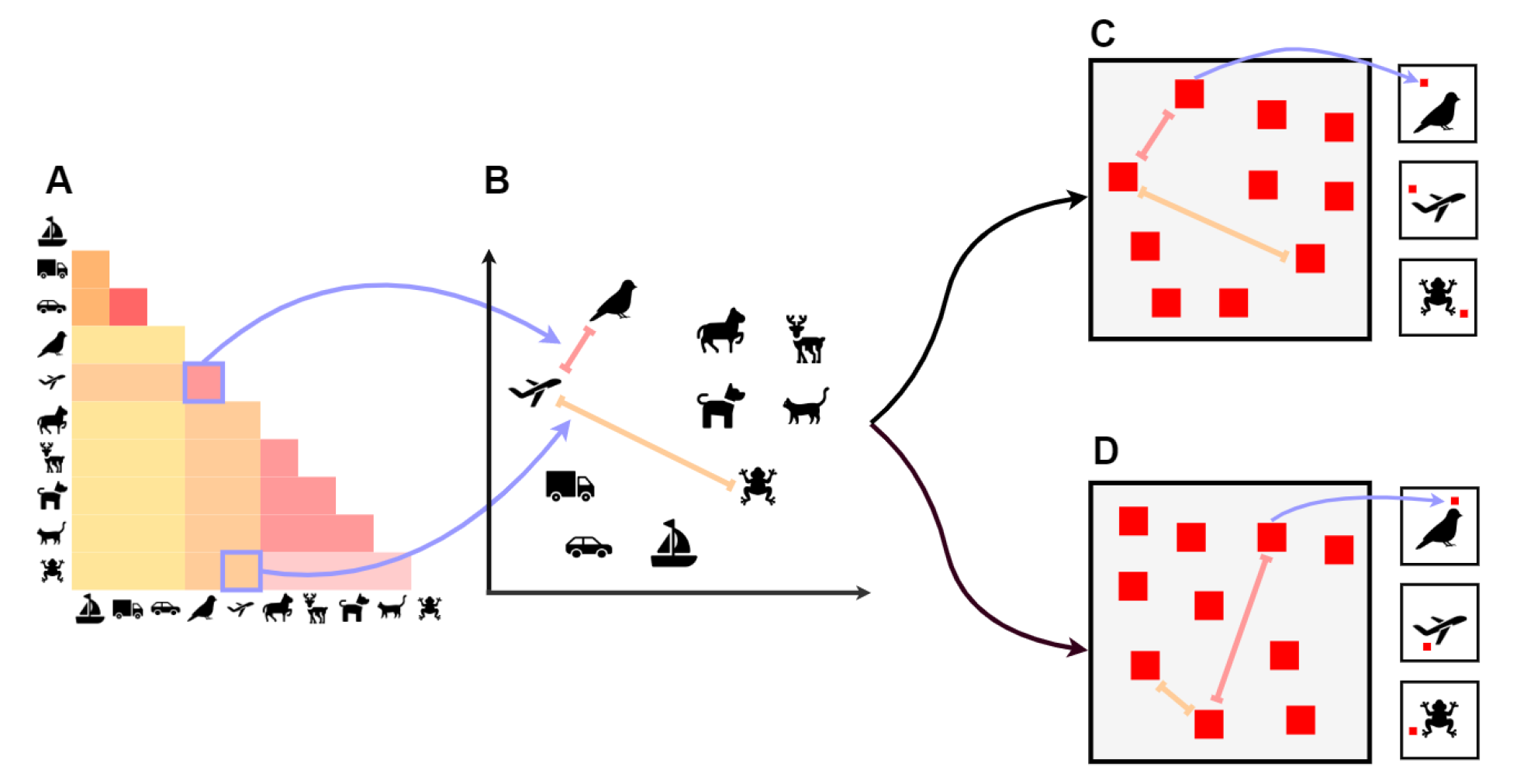
Study 2 confound placement. The representational geometry (Panel A and B) from the network trained on the unperturbed `CIFAR-10` images is used to determine the location of the single pixel confound (shown as a red patch here) for each category. In the ‘Positive’ condition (Panel C), we determined 10 locations in a 2D plane such that the distances between these locations were positively correlated to the representational geometry – illustrated here as the red patches in Panel C being in similar locations to category locations in Panel B. These 10 locations were then used to insert a single diagnostic – i.e., category-dependent – pixel in each image (Insets in Panel C). A similar procedure was also used to generate datasets where the confound was uncorrelated (Panel D) or negatively correlated (not shown here) with the representational geometry of the network.

The locations of these diagnostic pixels were chosen such that they were correlated to the corresponding representational distances between classes in Φ_1_. Our hypothesis was that if the representational distances in Φ_2_ preserve the physical distances of diagnostic pixels in input space, then this confound will end up mimicking the representational geometry of Φ_1_, even though the two systems use qualitatively different features for classification. Furthermore, we trained two more networks, Φ_3_ and Φ_4_, which were identical to Φ_2_, except these networks were trained on datasets where the location of the confound was uncorrelated (Φ_3_) or negatively correlated (Φ_4_) with the representational distances in Φ_1_ (see Fig 5 and Methods for details).

Classification accuracy (Fig 6 (left)) revealed that the network Φ_1_, trained on the unperturbed images, learned to classify these images and ignored the diagnostic pixel – that is, it’s performance was identical for the unperturbed and modified images. In contrast, networks Φ_2_ (positive), Φ_3_ (uncorrelated) and Φ_4_(negative) failed to classify the unperturbed images (performance was near chance) but learned to perfectly classify the modified images, showing that these networks develop qualitatively different representations compared to normally trained networks.

**Figure 6:**
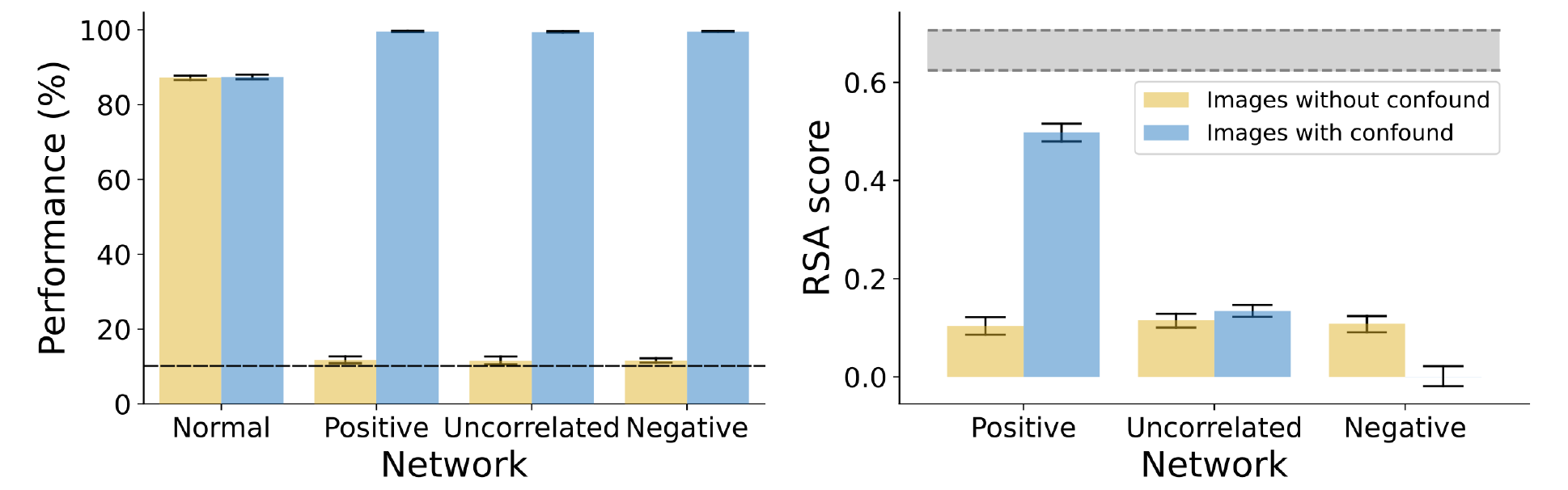
Study 2 results. *Left:* Performance of normally trained networks did not depend on whether classification was done on unperturbed `CIFAR-10` images or images with a single pixel confound (error bars represent 95% CI, the dashed line represents chance performance). All three networks trained on datasets with confounds could perfectly categorise the test images when they contained the confound (blue bars), but failed to achieve above-chance performance if the predictive pixel was not present (yellow bars). *Right:* The RSA score between the network trained on the unperturbed dataset and each of the networks trained on datasets with confounds. The three networks showed similar scores when tested on images without confounds, but vastly different RSA scores when tested on images with confounds. Networks in the Positive condition showed near ceiling scores (the shaded area represents noise ceiling) while networks in the Uncorrelated and Negative conditions showed much lower RSA.

Next we computed pairwise RSA scores between the representations at the last convo-lution layer of Φ_1_ and each of Φ_2_, Φ_3_ and Φ_4_ (Fig 6 (right)). When presented unperturbed test images, the Φ_2_, Φ_3_ and Φ_4_ networks all showed low RSA scores with the normally trained Φ_1_ network. However, when networks were presented with test images that in-cluded the predictive pixels, RSA varied depending on the geometry of pixel locations in the input space. When the geometry of pixel locations was positively correlated to the normally trained network, RSA scores approached ceiling (i.e., comparable to RSA scores between two normally trained networks). Networks trained on uncorrelated and negatively correlated pixel placements scored much lower.

These results mirror Study 1: we observed that it is possible for two networks (Φ_1_ and Φ_2_) to show highly correlated representational geometries even though these networks learn to classify images based on very different features. One may argue that this could be because the two networks could have learned similar representations at the final convolution layer of the DNN and it is the classifier that sits on top of this representation that leads to the behavioural differences between these networks. But if this was true, it would not explain why RSA scores diminish for the two other comparisons (with Φ_3_ and Φ_4_). This modulation of RSA-scores for different datasets suggests that, like in Study 1, the correlation in representational geometry is not because the two systems encode similar features of inputs, but because different features mimic each other in their representational geometries.

### Re-examining some influential findings

In Studies 1 and 2, we showed that it is possible for qualitatively different systems to end up with similar representational geometries. However, it may be argued that while this is possible in principle, it is unlikely in practice in real-world scenarios. In the following two studies, we consider real-world data from some recent influential experiments, recorded from both primate and human cortex. We show how RSA-scores can be driven by confounds in these real-world settings and how properties of training and test data may contribute to observed RSA-scores.

### Study 3: Neural activations in monkey IT cortex can show a high RSA-score with DNNs despite different encoding of input data

In our next study, we consider data from experiments comparing representational geometries between computa-tional models and macaque visual cortex [12, 30]. The experimental setup was similar to Study 2, though note that unlike Study 2, where both systems used the same archi-tecture and learning algorithm, this study considered two very different systems – one artificial (DNN) and the other biological (macaque IT cortex). We used the same set of images that were shown to macaques by Majaj et al. [31] and modified this dataset to superimpose a small diagnostic patch on each image. In the same manner as in Study 2 above, we constructed three different datasets, where the locations of these diagnostic patches were either positively correlated, uncorrelated or negatively correlated with the RDM of macaque activations. We then trained four CNNs. The first CNN was pre-trained on `ImageNet` and then fine-tuned on the unmodified dataset of images shown to the macaques. Previous research has shown that CNNs trained in this manner develop representations that mirror the representational geometry of neurons in primate inferior temporal (IT) cortex [12]. The other three networks were trained on the three modi-fied datasets and learned to entirely rely on the diagnostic patches (accuracy on images without the diagnostic patches was around chance).

Fig 7 (right) shows the correlation in representational geometry between the macaque IT activations and activations at the final convolution layer for each of these networks. The correlation with networks trained on the unmodified images is our baseline and shown as the gray band in Fig 7. Our first observation was that a CNN trained to rely on the di-agnostic patch can indeed achieve a high RSA score with macaque IT activations. In fact, the networks trained on patch locations that were positively correlated to the macaque RDM matched the RSA score of the CNNs trained on `ImageNet` and the unmodified dataset. This shows how two systems having very different architectures, encoding fun-damentally different features of inputs (single patch vs naturalistic features) can show a high correspondence in their representational geometries. We also observed that, like in Study 2, the RSA score depended on the clustering of data in the input space – when patches were placed in other locations (uncorrelated or negatively correlated to macaque RDMs) the RSA score became significantly lower.

**Figure 7:**
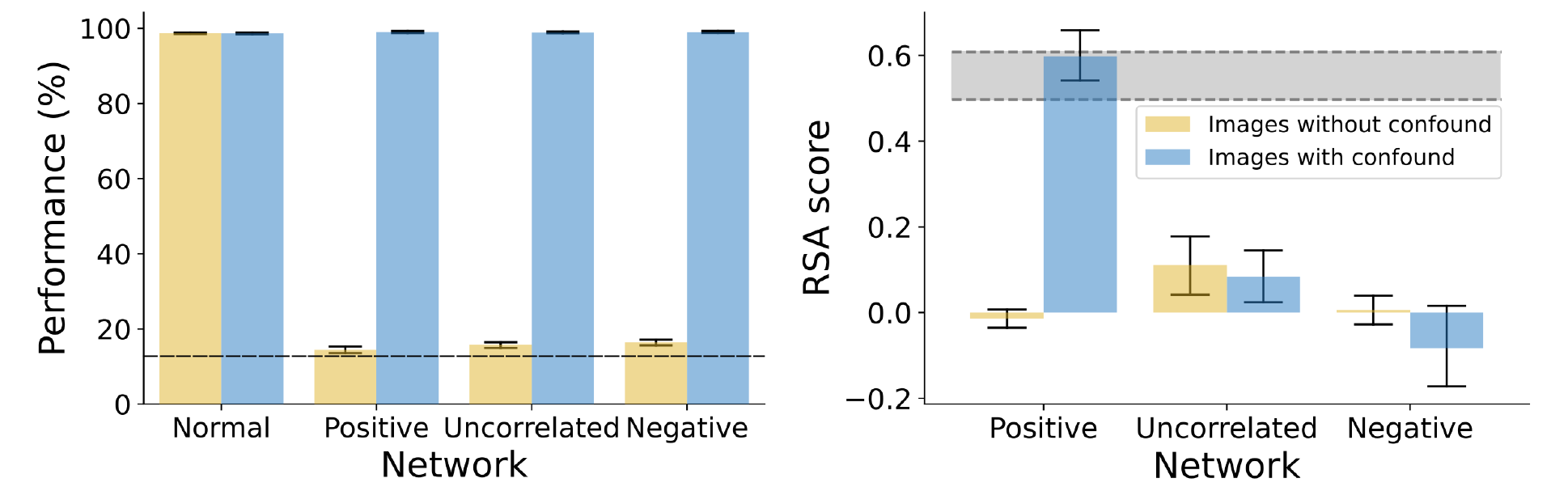
Study 3 results. *Left:* Classification Performance of the network trained on unperturbed images (Normal condition) did not depend on the presence or absence of the confound, while performance of networks trained with the confound (Positive, Uncorrelated and Negative conditions) highly depended on whether the confound was present (dashed line represents chance performance). *Right:* RSA-scores with macaque IT activations were low for all three conditions when images did not contain a confound (yellow bars). When images contained a confound (blue bars), the RSA-scores depended on the condition, matching the RSA-score of the normally trained network (grey band) in the Positive condition, but decreasing significantly in the Uncorrelated and Negative conditions. The grey band represents a 95% CI for the RSA-score between normally trained networks and macaque IT activations.

### Study 4: High RSA-scores may be driven by the structure of testing data

All the studies so far have used the same method to construct datasets with confounds – we established the representational geometry of one system (Φ_1_) and constructed datasets where the clustering of features (pixels) mirrored this geometry. However, it could be argued that confounds which cluster in this manner are unlikely in practice. For example, even if texture and shape exist as confounds in a dataset, the inter-category distances between textures are not necessarily similar to the inter-category distances between shape. However, categories in real-world datasets are usually hierarchically clustered into higher-level and lower-level categories. For example, in the `CIFAR-10` dataset, the Dogs and Cats (lower-level categories) are both animate (members of a common higher-level category) and Airplanes and Ships (lower-level categories) are both inanimate (members of a higher-level category). Due to this hierarchical structure, Dog and Cat images are likely to be closer to each other not only in their shape, but also their colour and texture (amongst other features) than they are to Airplane and Ship images. In our next simula-tion, we explore whether this hierarchical structure of categories can lead to a correlation in representational geometries between two systems that learn different feature encodings.

For this study, we selected a popular dataset used for comparing representational geometries in humans, macaques and deep learning models [13, 32]. This dataset consists of six categories which can be organised into a hierarchical structure shown in Fig 8. [6] showed a striking match in RDMs for response patterns elicited by these stimuli in human and macaque IT. For both humans and macaques, distances in response patterns were larger between the higher-level categories (animate and inanimate) than between the lower-level categories (e.g., between human bodies and human faces).

**Figure 8:**
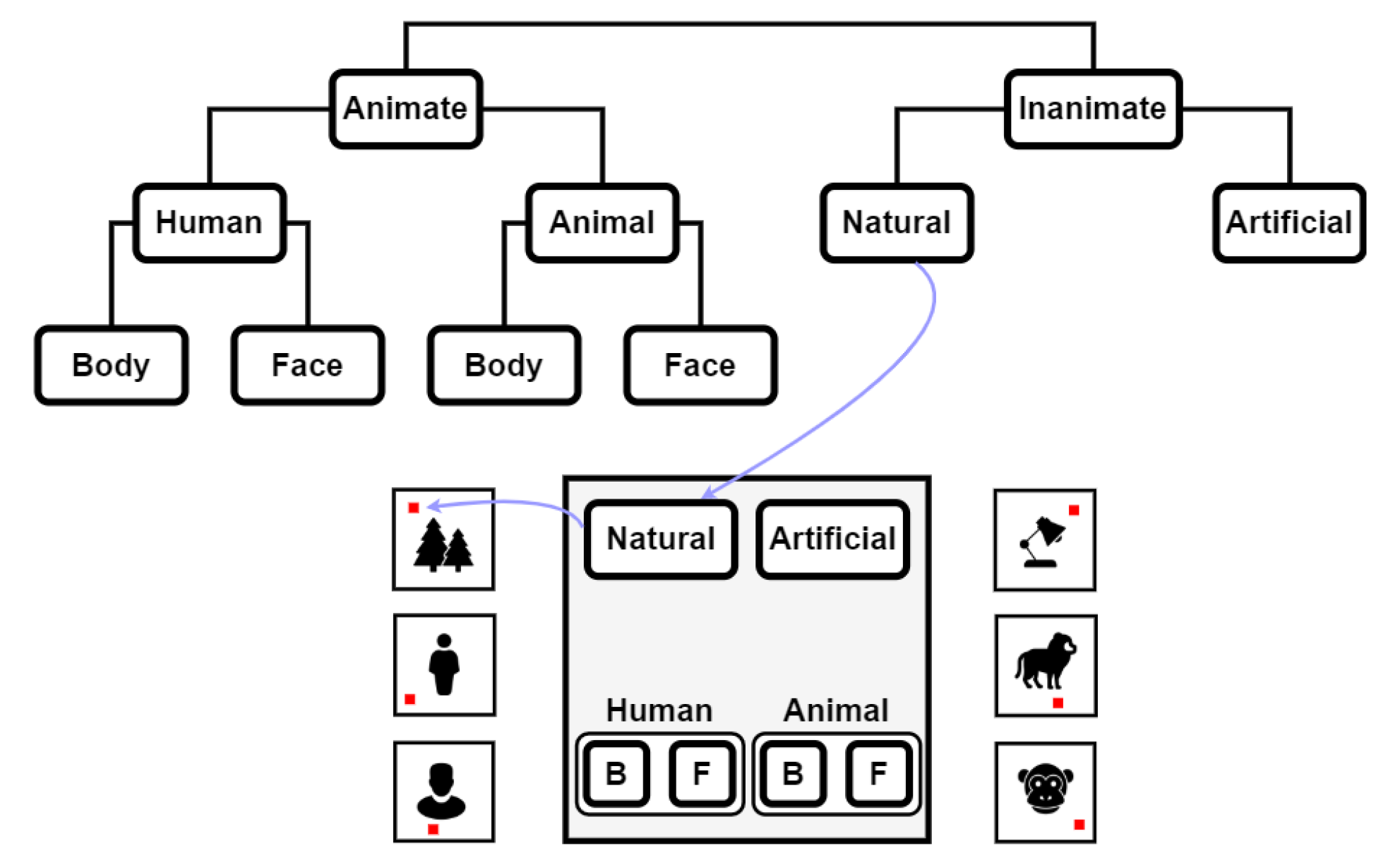
Exploiting intrinsic dataset hierarchy in order to place confounds. The top panel shows the hierarchical structure of categories in the dataset, which was used to place the single pixel confounds. The example at the bottom (middle) shows one such hierarchical placement scheme where the pixels for Inanimate images were closer to the top of the canvas while Animate images were closer to the bottom. Within the Animate images, the pixels for Humans and Animals were placed at the left and right, respectively, and the pixels for bodies (B) and faces (F) were clustered as shown.

We used a similar experimental paradigm to the above studies, where we trained networks to classify stimuli which included a single predictive pixel. But instead of using an RDM to compute the location of a diagnostic pixel, we used the hierarchical categorical structure. In the first modified version of the dataset, the location of the pixel was based on the hierarchical structure of categories in Fig 8 – predictive pixels for animate kinds were closer to each other than to inanimate kinds, and pixels for faces were closer to each other than to bodies, etc. One such configuration can be seen in Fig 8. In the second version, the predictive pixel was placed at a random location for each category (but, of course, at the same location for all images within each category). We call these conditions ‘Hierarchical’ and ‘Random’. [13] showed that the RDM of average response patterns elicited in the human IT cortex (Φ_1_) correlated with the RDM of a DNN trained on naturalistic images (Φ_2_). We explored how this compared to the correlation with the RDM of a network trained on the Hierarchical pixel placement (Φ_3_) and Random pixel placement (Φ_4_).

Results for this study are shown in Fig 9. We observed that representational geometry of a network trained on Hierarchically placed pixels (Φ_3_) was just as correlated to the representational geometry of human IT responses (Φ_1_) as a network trained on naturalistic images (Φ_2_). However, when the pixel locations for each category were randomly cho-sen, this correlation decreased significantly. These results suggest that any confound in the dataset (including texture, colour or low-level visual information) that has distances governed by the hierarchical clustering structure of the data could underlie the observed similarity in representational geometries between CNNs and human IT. More generally, these results show how it is plausible that many confounds present in popular datasets may underlie the observed similarity in representational geometries between two systems. The error of inferring a similarity in mechanism based on a high RSA score is not just possible but also probable.

**Figure 9:**
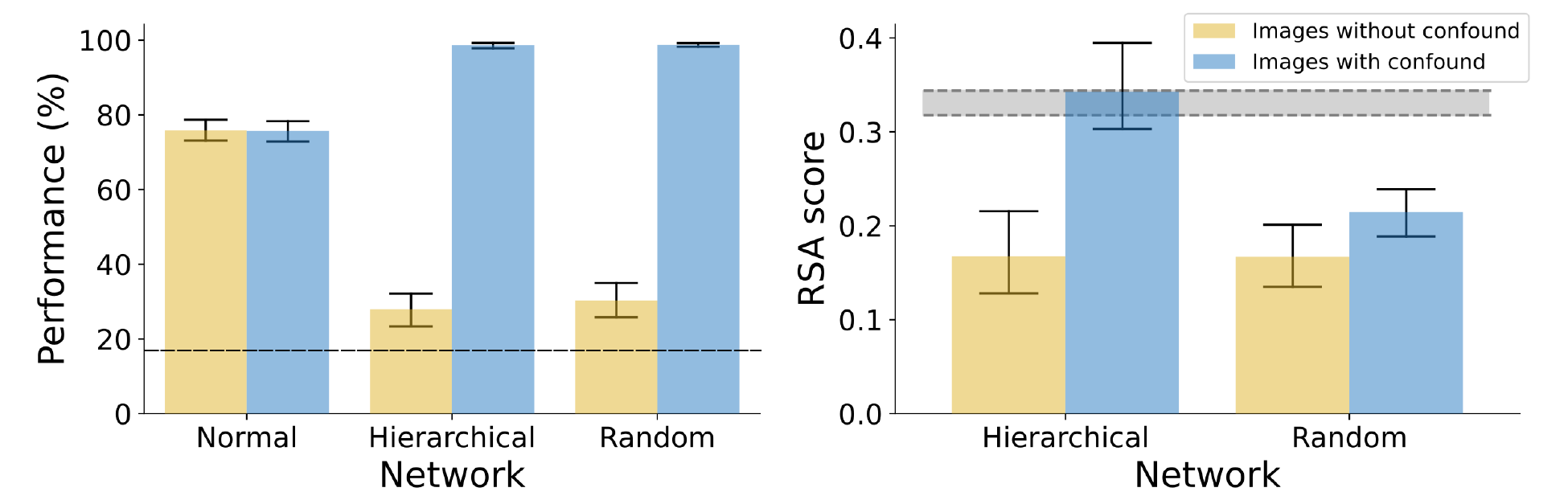
Study 4 results. *Left:* Performance of normally trained networks did not depend on whether the confound was present. Networks trained with the confound failed to classify stimuli without the confound (yellow bars) while achieving near perfect classification of stimuli with the confound present (blue bars, dashed line represents chance performance). *Right:* RSA with human IT activations reveals that, when the confound was present, the RSA-score for networks in the Hierarchical condition matched the RSA-score of normally trained network (gray band), while the RSA-score of the network in the Random condition was significantly lower. The grey band represents 95% CI for the RSA score between normally trained networks and human IT.

## Discussion

In four studies, we have illustrated a number of conditions under which it can be problem-atic to infer a similarity of representations between two systems based on a correlation in their representational geometries. In particular, we showed that two systems may transform their inputs through very different functions and encode very different features of inputs and yet have highly correlated representational geometries. Of course, one may acknowledge that a second-order isomorphism of activity patterns does *not* strictly imply that two systems are similar mechanistically but still assume that it is highly likely to be the case. That is, as a practical matter, a researcher may assume that RSA is a reliable method to compare complex systems representing high-dimensional inputs. However, our findings challenge this assumption. We show how a high RSA score between different systems can not only occur in a bare-bones simulation (Study 1), but also in practice, in high-dimensional systems operating on high-dimensional data (Studies 2–3). Furthermore, we show that the hierarchical structure of datasets frequently used to test similarity of representations lends itself to artificially inflated RSA scores because of second-order confounds present in the dataset (Study 4). Therefore, second-order confounds driving high RSA scores is not only possible but plausible.

### Significance and Implications

These results are critical for any researchers interested in using RSA for comparing mechanistic processes of complex systems. This includes research comparing information processing across species [6], across brain areas [9], between computational models [33] and between artificial intelligence models and brains [12–16]. Our findings are particularly relevant to an ongoing debate about similarity of information processing in Deep Neu-ral Networks and the mamillian visual cortex. Studies comparing Convolutional Neural Networks (CNNs) to the visual cortex present a set of contradictory findings. In some studies, researchers have observed a correlation between the geometry of activation patterns in a CNN trained to classify large datasets of images and the geometry of neural activation patterns within the macaque or human inferotemporal cortex, when both systems are stimulated using a set of naturalistic images [13]. Based on these observations of similarity (in RDMs), many researchers infer that CNNs provide insight into the mechanisms of information processing in the visual cortex [14, 15]. But other researchers have challenged this claim arguing that there are too many differences – from architectures and algorithms to tasks and environments used for learning – to meaningfully compare these systems, and indeed many behavioural results provide striking contrast between the performance of humans and CNNs [34–36]. Our results show how it is possible (and plausible) for these contradictory results to arise. That is, it is possible that CNNs are extracting very different features of naturalistic images and may even be transforming these images through very different functions compared to biological visual systems and yet end up with correlated representational geometries due to confounds present in datasets and their structural properties. It is important to emphasize that confounds are ubiquitous in datasets [29] leading DNNs to often classify images on the basis of short-cuts [37] and it unclear why confounds would not also drive high RSA scores. Furthermore, our results are consistent with with recent results such as Xu and Vaziri-Pashkam [38] who have reported that the correlation between representational geometries (previously reported in many studies) depends on the dataset used for testing. Still, whether this is actually happening will require finding the actual confound driving these effects, a non-trivial task in such high-dimensional data (see below under ‘Limitations’).

We would also like to emphasise that our results are not an indictment of RSA per se. Rather, our critique is aimed at problematic inferences that are frequently being drawn based on this statistical method when comparing complex systems. Like many statistical methods, Representation Similarity Analysis provides insight into data at a certain level of abstraction. There are many fruitful ways to use RSA, particularly by constructing theory-based representation dissimilarity matrices and comparing observed RDMs with these theory-driven RDMs. An example of this approach is outlined by Naselaris & Kay [18], who discuss how RDMs can be constructed based on hypotheses (such as whether the luminance of an image is driving observed differences between conditions) and compared with the RDMs of observed data. Used in this way, RSA is a useful tool for model-comparison, with different target RDMs corresponding to clearly defined hypotheses. The problem arises when the target RDM is generated by a complex system processing high-dimensional stimuli. In this setting, our results show that alternative hypotheses about mechanism are difficult to disambiguate based on RSA.

### Philosophical implications

But couldn’t a researcher take the view that representational geometry *is* representation and therefore, a strong correlation in representational geometries between two systems is sufficient to infer that the systems are representing the world in a similar manner? This question goes to the heart of an existing debate in philosophy, where philosophers distin-guish between the *externalist* and *holistic* views on mental representations [39]. According to the first view, the content of representations is determined by their relationship to enti-ties in the external world. This perspective is implicitly taken by most neuroscientists and psychologists, who are interested in comparing mechanisms underlying cognitive processes – that is, they are interested in the set of nested functions and algorithms responsible for transforming sensory input into a set of activations in the brain. From this perspective, our finding that high RSA scores can be obtained between systems that work in quali-tatively different ways poses a challenge to researchers using RSA to compare complex systems using high-dimensional data where a multitude of feature transformations could be driving the observed results.

Alternatively, a researcher may reject the externalist view and adopt the perspective that representations obtain their meaning based on how they are related to each other within each system, rather than based on their relationship to entities in the external world. That is, “representation *is* the representation of similarities” [40]. From this per-spective, as long as the two systems share the same relational distances between internal activations, one can validly infer that the two systems have similar representations. That is, a second-order isomorphism implies a similarity of representations, by definition. This view has been called *holism* in the philosophy of mind [39, 41] and is related to a similar idea of *meaning holism* in language, which is the idea that the meaning of a linguistic expression is determined by its relation to other expressions within a language [42, 43]. For example, Firth [44] (p. 11) writes: “you shall know a word by the company it keeps”. Similarly, Griffiths and Steyvers [45], and Griffiths, Steyvers, and Tenenbaum [46] have adopted meaning holism accounts of semantic representations in neural networks. More recently, Piantadosi and Hill [47] have argued that large language models capture impor-tant aspects of meaning and approximate human cognition providing one assumes meaning arises from the relations of states rather than an architecture or training. Even if a re-searcher were to adopt this holistic perspective on representations, our results should still be of interest to them as they show that the similarity between representational geome-tries can vary based on the visual stimulus that is used to compare them (the modulation effect). As all our studies show, representational geometries heavily depend on the dataset - e.g., a networks trained with a confound in the training set will have vastly different geometries when presented test data with and without the confound present. Addition-ally, our results show that adopting this view misses the information about differences in mechanistic processes that a psychologist or neuroscientist is frequently interested in, for instance, whether the visual system processes surface reflectance or shape in order to identify objects. Fodor and Lepore long ago criticized this philosophical stance [39, 48], and interestingly, this philosophical debate played an important part in the development of RSA (see section S1 of Supplementary Information). Unfortunately, this debate has largely been ignored by contemporary researchers who use RSA as a method to infer similarity of systems.

### Relation to existing research

A related point has been made by Kriegeskorte and Diedrichsen [22] and Kriegeskorte and Wei [49], who point out that two systems may have the same representational geometry, even if they have a different activity profile over neurons. In this sense, the geometry abstracts away the information about how information was distributed over a set of neurons. Kriegeskorte and Diedrichsen [22] equate this loss in information to “peeling a layer of an onion” – downstream decoders that are sensitive to the representational geometry rather than activity profiles over neuron populations can focus on difference in information as reflected by a change in geometry and be agnostic to how this information is distributed over a set of neurons. We agree that this invariance over activity profiles is indeed a useful property of representational geometries for downstream decoders. However, while abstracting over activity profiles may be useful, abstracting over stimulus properties loses an important piece of information when comparing representations across brain regions, individuals, species and between brains and computational models. Our studies show how two systems may appear similar based on their representational geometries in one circumstance (e.g. Fig 2B) but drastically different in another circumstance (Fig 2C). Furthermore, our results show how such second-order confounds can arise because of properties of transformations (Study 1) and structural properties of datasets used for testing (Study 4).

It is also important to note how our results differ from previous studies exploring limitations of RSA. Some of these studies have focused on the importance of how neural data is pre-processed. For example, Ramirez [50] found that pre-processing steps, such as centering (de-meaning) activation vectors may lead to incorrect inference about the representational geometry of activations. They demonstrated that subtracting the mean from activations could change the rank order of similarity between conditions. In turn, this could lead to clearly distinct RDMs becoming highly correlated and vice-versa. While this is an important methodological point, it is clearly distinct from the point we are making in this study. Indeed, the results here are agnostic of the data pre-processing steps and hold whether or not activations are centered.

Another set of studies have explored how the procedure of data collection can influence the results of RSA. For example, Henriksson et al. [51] and Cai et al. [52] demonstrated that RDMs measured based on fMRI data can be severely biased because of temporal and spatial correlations in neural activity. These authors have pointed out that if activity patterns from different brain regions are recorded during the same trial, the similarity estimates will be exaggerated due to correlated neural fluctuations in these regions. Sim-ilarly, neural activity is correlated over time, which means estimated similarity based on activity patterns from the same imaging run also introduces a strong bias in RDMs. These sources of bias are important to understand, but they can also be addressed by a more careful task design and analysis [52]. In contrast, the confounds that are highlighted in this study exist in the stimulus itself. Therefore, even if one were to completely mitigate the bias in estimating RDMs, the types of confounds we highlight in our work would still pose problems when drawing inferences from correlation in RDMs.

A third set of studies have highlighted the importance of choosing the correct distance metric when using RSA. For example, Ramirez [26] compared Euclidean distance with an angular metric (such as cosine similarity) and showed that the choice of distance metric can reveal different aspects of the same fMRI data. They argued that the Euclidean distance is particularly sensitive to the mean activity over a recorded voxel. Based on this analysis, Ramirez [26] suggested using an angular distance metric, especially when neural signal is aggregated over large number of neurons. Similarly, in another exhaustive study over distance measures, Bobadilla-Suarez et al. [25], evaluated neural similarity using various distance measures, including angle-based measures (cosine, Pearson, Spearman) and magnitude-based measures (Euclidean, Mahalanobis, Minkowski) and found that the choice of metric significantly influenced the measured similarity. They also found that there was no one metric that outperformed all others – rather, the preferred metric varied across different studies, but was consistent across brain regions within a study. The choice of distance metric is again a related but orthogonal issue to the one we highlight in this study. Representational geometry abstracts away information about stimulus features and how inputs are transformed. Our results demonstrate how different stimulus features and transformations of input stimulus can lead to the same representational geometry. This is integral to the nature of representational geometries, rather than a consequence of the distance metric used.

Of course, the problem of confounds in stimuli is not unique to RSA and will affect other statistical analyses, including multivariate regression methods such as MVP classification. A number of studies have pointed out this conceptual problem in the context of multivariate decoding, where authors have argued that successfully decoding a signal from a neural activation pattern is no guarantee that the signal is encoded by the brain or decoded by downstream processes [18–20, 53]. In response, researchers have adopted a variety of methods to deal with such confounds such as cross-validation [23], confound regression [24], counter-balancing data [54] and commonality analysis [55]. We couldn’t agree more with this direction of research and our study highlights two properties of confounds that makes it especially challenging to compare neural representations with those in complex models working on high-dimensional data. Firstly, these confounds are second-order – that is, they are not only category-correlated (as is the case for confounds highlighted for multivariate decoding), but also mimic the second-order similarity struc-ture of the variable of interest. Secondly, when using high-dimensional datasets (such as naturalistic images) and complex target models (such as DNNs) for testing, these con-founds are unknown to the experimenter and may be present in the entire dataset. This restricts the utility of existing methods, such as cross-validation and counter-balancing data, for dealing with these confounds, a point made by researchers employing these methods [24, 54]. In fact, we are unaware of any statistical methods that can completely eliminate confounds under these settings.

### Limitations and General Recommendations

Even though our findings demonstrate that second-order confounds are possible and plau-sible, they do not allow us to infer whether such confounds *are* present in existing datasets and driving the observed similarity in existing studies. An important research direction is to discover these confounds (or lack thereof) and determine the extent to which repre-sentations in a target model mimic the second-order relationship between neural repre-sentations in the visual cortex. One way to do this is to systematically eliminate possible confounds from datasets and check the extent to which this affects previously observed results. Of course, this is not straightforward to do in high-dimensional stimuli, such as naturalistic images, which consist of millions of features and combinatorial relation-ships between these features. Thus identifying confounds in such datasets remains a real challenge.

In closing, we describe our recommendations for practitioners who would like to use RSA for comparing complex systems based on high-dimensional data. First, since the intrinsic structure of datasets can artificially modulate RSA scores, researchers should compare systems on a wider variety of datasets and sampling schemes than currently done. Second, given that confounding features can lead to mimicked representational geometries, researchers should consider running additional controlled experiments that manipulate independent variables designed to test hypotheses to rule out this possibility when inferences hinge crucially on it. This point has recently been made by Bowers et al [36] in relation to testing the similarities of DNN and human vision. Similarly, the ‘controversial stimuli’ designed by Golan et al. [56] should also enable researchers to test representational geometries for stimuli where different models make contrasting predictions. Third, when studies are conducted to search for evidence of mechanistic similarity between two or more systems, researchers should use a wider range of complementary methods in order to increase robustness (e.g., RSA combined with neural predictivity [12], MVPC [7, 57], CCA [58], SVCCA [59], CKA [60]).

Lastly, perhaps the most important general recommendation we make is that researchers should acknowledge, procedurally and in writing, which inferences are afforded by the use of RSA, and what dissimilarities remain possible despite having observed a given pattern of RSA scores. To this end, we believe that general statements of similarity tend to obfuscate rather than accurately summarize any set of RSA-based results. Instead, we urge researchers using RSA (1) to justify the use of this method by theoretically motivated interest in representational geometry or otherwise consider other tools that best fit their goals, and (2) to state in precise terms that RSA scores reflect the similarity of representational geometries in particular, and generally avoid underspecified claims of similarity.

## Methods

### Dataset generation and training

All DNN simulations (Studies 2–4) were carried out using the `Pytorch` framework [61]. The model implementations were downloaded from the `torchvision` library. Networks trained on unperturbed datasets in all studies were pre-trained on `ImageNet` as were networks trained on modified datasets in Study 2. Networks trained on modified datasets in Studies 3 and 4 were randomly initialised. For the pre-trained models, their pre-trained weights were downloaded from `torchvision.models` subpackage.

### Study 1

Each dataset in Study 1 consists of 100 samples (50 in each cluster) drawn from two multivariate Gaussians, *N* (*x|µ,* **Σ**), where *µ* is a 2-dimensional vector and **Σ** is a 2 *×* 2 covariance matrix. In Fig 2A, the two Gaussians have means *µ***_1_** = (1, 8) and *µ***_2_** = (8, 1) and a covariance matrices 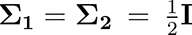, while in Fig 2B the Gaussians have means *µ***_1_** = (1, 1) and *µ***_2_** = (8, 8) and a covariance matrices **Σ_1_** = **I**, **Σ_2_** = 8**I**. All kernel matrices were computed using the `sklearn.metrics.pairwise` module of the `scikit-learn` Python package.

### Study 2

First, a VGG-16 deep convolutional neural network [62], pre-trained on the `ImageNet` dataset of naturalistic images, was trained to classify stimuli from the `CIFAR-10` dataset [63]. The `CIFAR-10` dataset includes 10 categories with 5000 training, and 1000 test images per category. The network was fine-tuned on `CIFAR-10` by replacing the classifier so that the final fully-connected layer reflected the correct number of target classes in `CIFAR-10` (10 for `CIFAR-10` as opposed to 1000 for `ImageNet`). Images were rescaled to a size of 224 *×* 224px and then the model learnt to minimise the cross-entropy error using the RMSprop optimizer with a mini-batch size of 64, learning rate of 10*^−^*^5^, and momentum of 0.9. All models were trained for 10 epochs, which were sufficient for convergence across all datasets.

Second, 100 random images from the test set for each category were sampled as input for the network and activations at the final convolutional layer extracted using the `THINGSVision` Python toolkit [64]. The same toolkit was used to generate a representational dissimilarity matrix (RDM) from the pattern of activations using `1-Pearson’s r` as the distance metric. The RDM was then averaged by calculating the median distance between each instance of one category with each instance of the others (e.g., the median distance between `Airplane` and `Ship` was the median of all pair-wise distances between activity patterns for airplane and ship stimuli). This resulted in a 10 *×* 10, category-level, RDM which reflected median between-category distances.

Third, three modified versions of the `CIFAR-10` datasets were created for the ‘Positive’, ‘Uncorrelated’ and ‘Negative’ conditions, respectively. In each dataset, we added one diagnostic pixel to each image, where the location of the pixel depended on the category (See Fig 5). The locations of these pixels were determined using the averaged RDM from the previous step. We call this the target RDM. In the ‘Positive’ condition, we wanted the distances between pixel placements to be positively correlated to the distances between categories in the target RDM. We achieved this by using an iterative algorithm that sampled pixel placements at random, calculated an RDM based on distances between the pixel placements and computed an RSA score (Spearman correlation) with the target RDM. Placements with a score above 0.70 were retained and further optimized (using small perturbations) to achieve an RSA-score over 0.90. The same procedure was also used to determine placements in the Uncorrelated (optimizing for a score close to 0) and Negatively correlated (optimizing for a negative score) conditions.

Finally, datasets were created using 10 different placements in each of the three condi-tions. Networks were trained for classification on these modified `CIFAR-10` datasets in the same manner as the VGG-16 network trained on the unperturbed version of the dataset (See Fig 4).

### Study 3

The procedure mirrored Study 2 with the main difference being that the target system was the macaque inferior temporal cortex. Neural data from two macaques, as well as the dataset were obtained from the Brain Score repository [30]. This dataset consists of 3200 images from 8 categories (animals, boats, cars, chairs, faces, fruits, planes, and tables), we computed an 8 *×* 8 averaged RDM based on macaque IT response patterns for stimuli in each category.

This averaged RDM was then used as the target RDM in the optimization procedure to determine locations of the confound (here, a white predictive patch of size 5 *×* 5 pixels) for each category. Using a patch instead of a single pixel was required in this dataset because of the structure and smaller size of the dataset (3200 images, rather than 50,000 images for `CIFAR-10`). In this smaller dataset, the networks struggle to learn based on a single pixel. However, increasing the size of the patch makes these patches more predictive and the networks are able to again learn entirely based on this confound (see results in Fig 6). In a manner similar to Study 2, this optimisation procedure was used to construct three datasets, where the confound’s placement was positively correlated, uncorrelated or negatively correlated with the category distances in the target RDM.

Finally, each dataset was split into 75% training (2432 images) and 25% test sets (768 images) before VGG-16 networks were trained on the unperturbed and modified datasets in the same manner as in Study 2. One difference between Studies 2 and 3 was that here the networks in the Positive, Uncorrelated and Negative conditions were trained from scratch, i.e., not pre-trained on `ImageNet`. This was done because we wanted to make sure that the network in the Normal condition (trained on `ImageNet`) and the networks in the Positive, Uncorrelated and Negative conditions encoded fundamentally different features of their inputs – i.e., there were no `ImageNet`-related features encoded by representations Φ_2_, Φ_3_ and Φ_4_ that were responsible for the similarity in representational geometries between these representations and the representations in macaque IT cortex.

### Study 4

The target system in this study was human IT cortex. The human RDM and dataset were obtained from [6]. Rather than calculating pixel placements based on the human RDM, the hierarchical structure of the dataset was used to place the pixels manually. The dataset consists of 910 images from 6 categories: human bodies, human faces, animal bodies, animal faces, artificial inanimate objects and natural inanimate objects. These low-level categories can be organised into the hierarchical structure shown in Fig 8. Predictive pixels were manually placed so that the distance between pixels for Animate kinds were closer together than they were to Inanimate kinds and that faces were closer together than bodies. This can be done in many different ways, so we created five different datasets, with five possible arrangements of predictive pixels. Results in the Hieararchical condition (Fig 9) are averaged over these five datasets. Placements for the Random condition were done similarly, except that the locations were selected randomly.

Networks were then trained on a 6-way classification task (818 training images and 92 test images) in a similar manner to the previous studies. As in Study 3, networks trained on the modified datasets (both Hierarchical and Random conditions) were not pre-trained on `ImageNet`.

### RDM and RSA computation

For Studies 2-4 all image-level RDMs were calculated using 1 *− r* as the distance measure. RSA scores were computed as the Spearman rank correlation between RDMs.

In Study 2, a curated set of test images was selected due to the extreme heterogeneity of the `CIFAR-10` dataset (low activation pattern similarity between instances of the same category). This was done by selecting 5 images per category which maximally correlated with the averaged activation pattern for the category. Since `CIFAR-10` consists of 10 categories, the RSA-scores in Study 2 were computed using RDMs of size 50 *×* 50.

In Study 3, the dataset consisted of 3200 images belonging to 8 categories. We first calculated a full 3200 *×* 3200 RDM using the entire set of stimuli. An averaged, category-level, 8 *×* 8 RDM was then calculated using median distances between categories (in a manner similar to that described for Study 2 in the Section ‘Dataset generation and training’). This 8 *×* 8 RDM was used to determine the RSA-scores. We also obtained qualitatively similar results using the full 3200 *×* 3200 RDMs. These results can be found in the S2 section of Supplementary Information.

In Study 4, the dataset consisted of 818 training images and 92 test images. Kriegeskorte et al. [6] used these images to obtain a 92*×*92 RDM to compare representations between human and macaque IT cortex. Here we computed a similar 92 *×* 92 RDM for networks trained in the Normal, Hierarchical and Random training conditions, which were then compared with the 92 *×* 92 RDM from human IT cortex to obtain RSA-scores for each condition.

### Testing

In Study 2, we used a 4 *×* 2 design to measure classification performance for networks in all four conditions (Normal, Postive, Uncorrelated and Negative) on both unperturbed images and modified images. We computed six RSA-scores: three pairs of networks – Normal-Positive, Normal-Uncorrelated and Normal-Negative – and two types of inputs – unperturbed and modified test images. The noise ceiling (grey band in Fig 6) was determined in the standard way as described in [65] and represents the expected range of the highest possible RSA score with the target system (network trained on the unperturbed dataset).

In Study 3, performance was estimated in the same manner as in Study 2 (using a 4*×*2 design), but RSA-scores were computed between RDMs from macaque IT activations and the four types of networks – i.e. for the pairs Macaque-Normal, Macaque-Positive, Macaque-Uncorrelated and Macaque-Negative. And like in Study 2, we determined each of these RSA-scores for both unperturbed and modified test images as inputs to the networks.

In Study 4, performance and RSA were computed in the same manner as in Studyn 3, except that the target RDM for RSA computation came from activations in human IT cortex and the networks were trained in one of three conditions: Normal, Hierarchical and Random.

### Data analysis

Performance and RSA scores were compared by running analyses of variance and Tukey HSD post-hoc tests. In Study 2 and 3, performance differences were tested by running a 4 (type of training) by 2 (type of dataset) mixed ANOVAs. In, Study 4, the differences were tested by running a 3×2 mixed ANOVA.

RSA scores with the target system between networks in various conditions were com-pared by running 3×2 ANOVAs in Studies 2 and 3, and a 2×2 ANOVA in Study 4. We observed that RSA-scores were highly dependent on both the way the networks were trained and also the test images used to elicit response activations.

For a detailed overview of the statistical analyses and results, see section S3 of the Supplementary Information.

## Data Availability

Confound placement coordinates (Studies 2-4), unperturbed datasets (Studies 3 and 4), macaque activation patterns and RDMs (Study 3) and human RDM (Study 4) are avail-able at OSF.

## Acknowledgments

This project has received funding from the European Research Council (ERC) under the European Union’s Horizon 2020 research and innovation programme (grant agreement No 741134)

## Footnotes

1 It is not our intention to pick on or criticise these authors here. We have inserted these quotes to point out the broad understanding of this method in the field. Many authors are indeed careful in stating that the term ‘similarity in representations’ is used as a shorthand for a ‘similarity in representational geometries’. Nevertheless, readers are also invited to accept that different systems show similar repre-sentational geometries because it is likely that they also use similar mechanisms to transform sensory information into latent representations, or they use similar (downstream) mechanisms to decode these latent representations e.g. [22]. But how safe are these assumptions?

