## Supplementary Information for "Obstacles to inferring mechanistic similarity using Representational Similarity Analysis"

### S1. Brief history of RSA

In the 1990s there was an important debate taking place on how to compare the mental representations of two individuals. On one side of this debate was Paul Churchland. Inspired by the success of connectionist models, Churchland argued that the brain represents reality as a pattern of activations over it's network of neurons [1]. This pattern of activation can be seen as a position in the brain's (high-dimensional) state-space. So, Churchland argued that one could compare how two individuals represent an object by comparing the corresponding positions in each individual's state-space. On the other side of the debate were Jerry Fodor and Ernie Lepore [2]. They pointed out that a problem with Churchland's proposal was that it "offers no robust account of content identity" (p 147). On Churchland's account, they argued, two mental representations have the same meaning only if they are embedded in identical state-spaces. This condition was highly unlikely to be satisfied in practice, given that no two brains have either the same number or connectivity of neurons and no two individuals have exactly the same experiences.

A possible solution to this problem of comparing representations across state-spaces of different dimensions was proposed by Laasko and Cottrell [3], who were investigating whether different neural networks, trained on the same data, represented an input stimulus in a similar manner. A direct comparison of activations across networks was not possible due to the difference in the number of units. To overcome this problem, [3] devised a method that compared encodings based on their *relative* positions in state-space. That is, based on a second-order isomorphism. They argued that two networks could be said to represent a concept in a similar manner if both networks partitioned their activation space (amongst concepts) in a similar manner – that is, if the activation spaces in both systems had a similar *geometry*. [3] conducted a series of experiments with neural networks, showing that neural networks with different sensory encodings and different number of hidden units nevertheless partitioned their activation space in a similar manner, leading them to conclude that these networks learned similar internal representations.

Churchland [4] saw Laasko and Cottrell's method as a decisive response to Fodor & Lepore's scepticism. He argued that, using Laasko and Cottrell's method, one could use the state-space approach to compare representations across individuals, even individuals that had different dimensions of their representational spaces. All one needed to do was to replace the requirement of "content identity" with "content similarity". That is, instead of comparing absolute positions of representations, one could simply compare how representations were organised *relative* to each other within each representational space.

However, Fodor & Lepore [5] argued that Churchland's reply was, in fact, "an egregious *ignoratio elenchi*" (p. 382). The problem was *not*, they argued, that one couldn't find the right metric to measure similarity across vector spaces of different dimensions. Rather, it was the fact that Churchland (and Laasko & Cottrell [3]) were interested in a *semantic* similarity – i.e., they wanted to compare whether representations had the same meaning in the two systems. Fodor & Lepore [5] argued that this problem of semantic similarity was intractable because similarity of concepts across systems of different dimensions is undefined. Consider the concept of a 'dog'. Let's say one person's representational space has a dimension of 'loyalty', while the other person's representational space does not. There is no principled answer for how similar the representation of 'dog' should be for these two individuals as it depends on how the dimension of 'loyalty' is weighted in the concept of 'dog'. And the relative weight of dimensions can differ for different concepts and circumstances. Moreover, Fodor & Lepore [5] argued that even identical representational geometries could *mean* very different things. For example, one individual may represent a dog along the dimensions of 'size' and 'speed' as being small (sized) and medium (speed).

Another individual may represent a dog along the dimensions of 'usefulness' and 'furriness' as being of small (usefulness) and medium (furriness). Even if the concept of a dog occupies a similar position in both state-spaces (small, medium) the two individuals clearly represent dogs differently.

Representation Similarity Analysis is an evolution of Laasko and Cottrell's method for comparing representations across systems. It retains its core principle of comparing representations based on their relative locations within each system's state-space. In addition, it formalises the ideas of similarity of representations within and across systems [6]. Like Laasko and Cottrell's method, a representation is usually coded as a vector of activation over some units (in a neural network or the brain). However, it could also be a behavioural measure, such as similarity judgments or even measures like accuracy or response times. We believe that many of the objections levelled by Fodor & Lepore against Churchland's idea of comparing systems based on relative positions in state-space also hold for representation similarity analysis. For example, Fodor & Lepore's point that similar state-space representations could mean different things can also be extended to RSA and in the main text we show how different systems with same representational geometries can, in fact, be encoding very different properties of sensory stimuli. From an externalist's perspective, activations within these systems *mean* very different things and yet have very comparable state-space representations (i.e. geometries). The only way to argue that concepts have a similar meaning in systems with similar representational geometries is to adopt a holistic perspective on representations. And as Fodor & Lepore [5] argued, and we discuss in the main text, adopting this perspective comes with its own set of problems.

### S2. Study 3 - Image-level RSA

Study 3 showed how RSA between networks sensitive to confounds and macaque inferior temporal cortex representational geometry can match the RSA score achieved by networks pretrained on naturalistic images and then fine-tuned on an unperturbed dataset. In the main paper we present category-level RSA scores – computed by first caluclating median distances between all instances of each category with all instances of each other category to get 8x8 RDMs which are then entered into RSA. Here, we present RSA scores without averaging. Activation patterns for each of the 3200 images in the dataset are used to calculate 3200x3200 RDMs which are used to compute RSA scores.

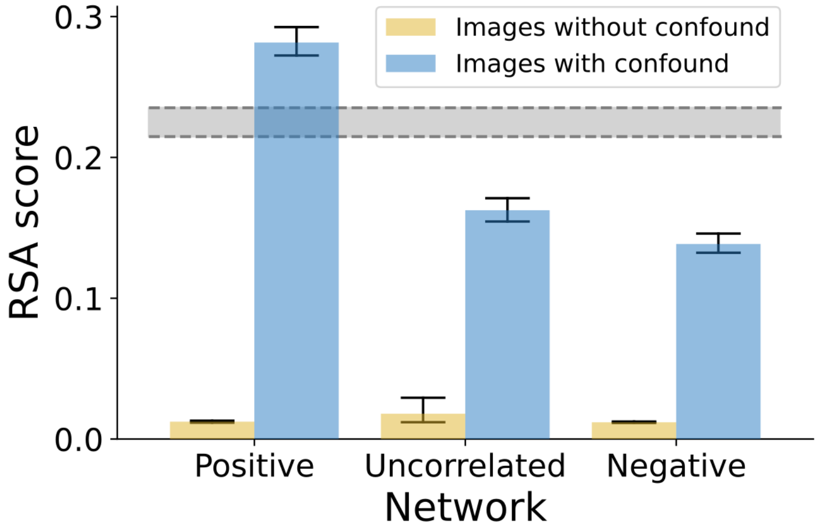


**Fig 1. Image-level RSA scores from Study 3.** RSA-scores with macaque IT activations were low for all three conditions when images did not contain a confound (yellow bars). When images contained a confound (blue bars), the RSA-scores depended on the condition, even exceeding the RSA-score of the normally trained network (grey band) in the Positive condition, but decreasing significantly in the Uncorrelated and Negative conditions. The grey band represents a 95% CI for the RSA-score between normally trained networks and macaque IT activations.

### S3. Statistical analyses

In this section we provide more detailed statistical analyses for Studies 2-4.

#### Study 2

In order to test for differences in performance (Figure 5, left panel in the main paper), a 4 (normally trained/positive/uncorrelated/negative) by 2 (dataset with/without confound) mixed analysis of variance (ANOVA) was conducted. The finding was a significant interaction effect ($F(3, 36) = 12256.10, p < .001, \eta_p^2 = .99$). Tukey HSD post-hoc comparisons revealed that performance in the positive, uncorrelated and negative conditions was significantly better on datasets which included the confounds (all $p < .001$) while the normally trained networks performed equally well on both datasets with and without the confound ($p = .99$). This shows that networks trained on datasets with confounds learned to classify based on the predictive confounding feature (single pixel) and ignored other features in the dataset (failing to classify if the confound is not present) while the normally trained networks remain unaffected by the presence or absence of the confounding feature.

Differences in RSA scores (Figure 5, right panel in the main paper) were tested by conducting a 3 (positive/uncorrelated/negative) by 2 (dataset with/without the confound) mixed ANOVA. The key findings was a significant interaction effect ($F(2, 297) = 289.27, p < .001, \eta_p^2 = .66$). Post-hoc comparisons revealed that there were no differences between the networks in RSA scores with normally trained networks when images without the confound were used as input (all $p > .954$). On the other hand, for images which contained the confound, networks in the positive condition achieved a significantly higher RSA score than both networks in the uncorrelated and negative conditions ($p < .001$), at the same time, networks in the uncorrelated condition achieved significantly higher RSA scores than networks in the negative condition ($p < .001$). This indicates a very strong modulation effect of RSA scores - depending on the relation between the representational geometry of the confounding feature exploited by these networks, RSA scores with normally trained networks can vary from high to low when the confound is present, but are consistently low when there is no confound in the test stimuli.

#### Study 3

The same analytical approach was taken as in Study 2, performance (Figure 6, left panel in the paper) was analyzed by conducting a 4 (normal/positive/uncorrelated/negative) by 2 (dataset with/without confound) mixed ANOVA. Again, the key finding was an interaction effect ($F(3, 51) = 8086.60, p < .001, \eta_p^2 = .99$). Post-hoc comparisons revealed that performance in the positive, uncorrelated and negative conditions was significantly better on datasets which included the confounds (all $p < .001$) while the normally trained networks performed equally well on both datasets with and without the confound ($p > .99$).

RSA scores (Figure 6, right panel in the main paper) were analyzed by conducting a 3 (positive/uncorrelated/negative) by 2 (dataset with/without confound) mixed ANOVA. The key result being a significant interaction effect ($F(2, 42) = 122.46, p < .001, \eta_p^2 = .85$). Post-hoc comparisons revealed that there were no differences between the networks in RSA scores with normally trained networks when images without the confound were used as input (all $p > .071$). However, for images with the confound present, networks in the positive condition achieve a significantly higher RSA score with macaque IT when compared to networks in the uncorrelated and negative conditions (all $p < .001$). Networks in the uncorrelated condition achieve higher RSA scores than networks in the negative condition ($p = .005$). Finally, it is worth emphasizing that networks in the positive condition match RSA scores with macaque IT achieved by networks pretrained on naturalistic images and then finetuned on the dataset without confounds ($t(23) = 0.89$, $p = .384$) when the confound is present in the dataset.

#### Study 4

For this simulation, performance differences between conditions (Figure 8, left panel in the main paper) were tested by conducting a 3 (normal/hierarchical/random) by 2 (dataset with/without confound) mixed ANOVA. As in previous studies, the eky result was a significant interaction effect ($F(2, 42) = 407.61, p < .001, \eta_p^2 = .95$). Post-hoc comparisons revealed that performance in the hierarchical and random conditions was significantly better on datasets which included the confounds (all $p < .001$) while the normally trained networks performed equally well on both datasets with and without the confound ($p > .99$).

RSA scores with human IT (Figure 8, right panel in the main paper) were analyzed by conducting a 2 (hierarchical/random) by 2 (dataset with/without) mixed ANOVA. The interaction effect was significant ($F(1, 28) = 8.46, p = .007, \eta_p^2 = .23$). Follow-up comparisons show that there was no difference between networks in the hierarchical and radnom conditions when the dataset did not contain the confound ($p > 99$), but networks in the hierarchical condition achieved significantly higher RSA scores when the dataset did contain the confound ($p < .001$). Again, it is worth emphasizing that networks in the hierarchical condition match RSA scores with human IT achieved by networks pretrained on naturalistic images and then finetuned on the dataset without confounds ($t(28) = 0.46$, $p = .647$) when the confound is present in the dataset.
